# Tributyltin Protects Against Ovariectomy-Induced Trabecular Bone Loss in C57BL/6J Mice with an Attenuated Effect in High Fat Fed Mice

**DOI:** 10.1101/2021.04.28.441846

**Authors:** Rachel Freid, Amira I Hussein, Jennifer J Schlezinger

**Author notes:** **Corresponding author and address for reprint requests:** Jennifer J. Schlezinger, Ph.D., Boston University School of Public Health Dept. of Environmental Health, 715 Albany Street, R-405 Boston, MA 02118. **Availability of Data and Material:** The authors will provide data upon request. **Code Availability:** Not applicable. **Authors Contributions:** J. J. S. designed the study, contributed to the experimental work and prepared the first draft of the paper. She is guarantor. R. F. and A. I. H. contributed to the experimental work and analyses. All authors revised the paper critically for intellectual content and approved the final version. All authors agree to be accountable for the work and to ensure that any questions relating to the accuracy and integrity of the paper are investigated and properly resolved. **Ethics Approval:** All animal studies were approved by the Institutional Animal Care and Use Committee at Boston University and performed in an American Association for the Accreditation of Laboratory Animal Care accredited facility (Animal Welfare Assurance Number: A3316-01).

## Abstract

Risk factors for poor bone quality include estrogen loss at menopause, a high fat diet and exposures to drugs/chemicals that activate peroxisome proliferator activated receptor gamma (PPARγ). We observed that the PPARγ and retinoid X receptor dual ligand, tributyltin (TBT), repressed periosteal bone formation but enhanced trabecular bone formation in female C57BL6/J mice. Here, we examined the interaction of diet, ovariectomy (OVX) and TBT exposure on bone structure. C57BL/6J mice underwent either sham surgery or OVX at 10 weeks of age. At 12 weeks of age, they were placed on a low (10% kcal) or high (45% kcal) fat, sucrose-matched diet and treated with Vh or TBT (1 or 5 mg/kg) for 14 weeks. OVX increased body weight gain in mice on either diet. TBT enhanced body weight gain in intact mice fed a high fat diet, but decreased weight gain in OVX mice. Elemental tin concentrations increased dose-dependently in bone. TBT had marginal effects on cortical and trabecular bone in intact mice fed a low- or high-fat diet. OVX caused a reduction in cortical and trabecular bone, regardless of diet. In high-fat fed OVX mice, TBT further reduced cortical thickness, bone area and total area. Interestingly, TBT protected against OVX-induced trabecular bone loss in low fat fed mice. The protective effect of TBT was nullified by the high fat diet and accompanied by a significant decrease in serum bone formation markers. Our novel observations will provide new information on basic bone biology, potential therapeutic targets and toxicological pathways.

## Introduction

Osteoporosis has been likened to “obesity of the bone” [1], as it results, in part, from the preferential differentiation of multipotent marrow stem cells (MSC) into adipocytes, at the expense of osteoblast differentiation [2]. The mutual-exclusivity of the adipogenesis and osteogenesis pathways most likely results from crosstalk between the major transcriptional regulators of these pathways, peroxisome proliferator activated receptor γ (PPARγ; adipogenesis) and Runx2 (osteogenesis)[3]. PPARγ activation induces differentiation along the adipocyte lineage and suppresses osteoblast differentiation [3, 4]. In osteocytes, PPARγ regulates the expression of sclerostin [5]. In osteoclasts, PPARγ is essential for differentiation via support of RANKL signaling [6, 7]. The importance of PPARγ in maintaining bone homeostasis is indicated by the fact that reduced PPARγ expression is associated with increased bone mass [3]. Treatment with therapeutic PPARγ agonists (such as rosiglitazone and pioglitazone used to treat type 2 diabetes) can exacerbate osteoporotic pathology particularly in post-menopausal women [8, 9].

Estrogen deficiency and consumption of a high fat and sugar, Western type diet are thought to contribute to osteoporosis risk in women [10]. It has been recognized for some time that loss of estrogen increases bone turnover [11]; estrogen-dependent control of bone remodeling is complex and results from the concerted interactions between both osteoclasts as well as osteoblasts and osteocytes (as reviewed in [12]). Either overexpression of PPARγ in osteoblasts or treatment with rosiglitazone significantly accelerates ovariectomy-induced bone loss [13, 14]. Consumption of a high fat diet increases expression of PPARγ in bone, and this is accompanied by loss of trabecular bone [15, 16]. While cortical bone mineral density has been shown to increase with high fat diet, bone quality decreases and brittleness increases [17]. The impact of a high fat diet on bone density and quality may be a result of both suppression of osteoblast differentiation and activation of bone resorption [18, 19].

Tributyltin (TBT) is a unique tool in that it is a dual agonist of PPARγ and its heterodimerization partner, the retinoid X receptors (RXR). TBT activates PPARγ:RXR and LXR:RXR heterodimers, as well as RXR homodimers [20, 21]. *In vitro*, TBT suppresses osteogenesis [22–25]. In young, female C57BL/6 mice, exposure to TBT reduces periosteal bone formation but increases trabecular bone formation [26]. This is consistent with observations in young female rats in which TBT induced an increase in whole body bone mineral density [27]. In young male rats, TBT only reduced bone mineral density in the diaphysis, and not the metaphysis, of the femur; the diaphysis contains substantially less trabecular bone than the metaphysis [28]. We hypothesized that the difference in effect in cortical versus trabecular bone results from the relative importance of osteoblast activity. In primary bone marrow macrophage cultures, TBT marginally inhibits the number of osteoclasts that differentiate but significantly suppresses expression of osteoclast markers *Nfatc1*, *Acp5* and *Ctsk* and resorptive activity [26, 29]. Gene expression studies revealed that TBT activated RXR- and LXR-dependent pathways *in vivo* and in osteoclast cultures. Both LXR and RXR have been proposed as targets for interventions to combat low bone density, given their roles in moderating osteoclast differentiation (as reviewed in [30, 31]).

Here, we investigated the effect of a unique chemical, TBT, on bone in the context of diet and ovary status in a C57BL/6J mouse model. The results show that TBT can protect trabecular bone from resorption stimulated ovariectomy, while having minimal effect on the cortical compartment. However, a high fat diet is shown to mitigate these potentially bone-protecting effects.

## Materials and Methods

### Materials

TBT chloride (T50202) was from Sigma-Aldrich (St. Louis, MO). All other reagents were from Thermo Fisher Scientific (Waltham, MA), unless noted.

### In vivo exposure

All animal studies were approved by the Institutional Animal Care and Use Committee at Boston University and performed in an American Association for the Accreditation of Laboratory Animal Care accredited facility (Animal Welfare Assurance Number: A3316-01). Exposures were carried out in 4 replicated experiments conducted over a 1 year and 9-month period. Female C57BL/6J mice (stock #:000664, RRID:IMSR_JAX:000664) that had either undergone sham surgery or ovariectomy at 10 weeks of age were purchased from Jackson Laboratories (Bar Harbor, ME). Beginning at 12 weeks of age, the mice were orally gavaged 3 times per week (Monday, Wednesday, Friday) for 14 weeks with vehicle (sesame oil, 5 μl/g), or TBT (1 or 5 mg/kg dissolved in sesame oil). Mice received either a low fat (10% fat as soybean oil and lard, 17% sucrose, D12450H, Research Diets, New Brunswick, NJ) or high fat diet (45% fat as soybean oil and lard, 17% sucrose, D12451, Research Diets). Mice were euthanized one day after the last dosing. Prior to euthanasia, mice were fasted overnight. Serum was collected at euthanasia and analyzed by ELISA for PINP (Rat/Mouse PINP EIA (AC-33F1), Immunodiagnostic Systems, Tyne &Wear, UK), Trap5b (Mouse Trap Assay (SB-TR103), IDS), and CTX (RatLaps EIA (AC-06F1), IDS). Livers were collected and analyzed for total triacylglyceride (TG) content as previously described [32]. Parametrial adipose tissues were collected and weighed. Humeri were collected and frozen at −20°C for ICP-MS analysis of tin content (Chemical Solutions Limited, Harrisburg, PA). Bone marrow was flushed from the marrow cavity of left femurs and left tibiae were collected; each were were flash frozen in liquid nitrogen and stored at −80°C until RNA was isolated. Right femurs were collected, wrapped in PBS soaked gauze, and frozen at −20°C until analyzed by micro-computed tomography. Right tibiae were collected and fixed in 4% paraformaldehyde.

### Micro-Computed Tomography (micro-CT)

Sample analyses were performed according to guidelines outlined in Bouxsein et al. (2010) [33]. Samples were scanned using a Scanco micro-CT 40 system (Scanco Medical; Brütisellen, Switzerland) using power, current, and integration time of 70 kVP, 113 μA, and 200 ms, respectively. Gaussian filtering (sigma = 0.8, support = 1) was used for reducing background noise. To analyze trabecular bone in the distal femur, the distal metaphysis was scanned at a nominal resolution of 6 μm/voxel, and the region of interest analyzed began at 0.03 mm proximal to the growth plate and extended 0.9 mm proximally. The trabecular compartment was manually segmented from the cortical shell. Cortical bone was analyzed from 12 μm/voxel scans of a 0.6 mm-tall region at the mid-diaphysis. For trabecular bone, treatment-specific thresholding was used (Vh = 438.5 mg HA/cm^3^; TBT = 455.2 mg HA/cm^3^) with thresholds determined by an iterative method (Scanco Medical); cortical bone was evaluated at a global threshold of 502 mg HA/cm^3^. Tissue mineral density was calculated with the aid of a standard curve obtained from a scan of a hydroxyapatite phantom consisting of five different mineral densities.

### Histology

Following 7 days of fixation at 4°C, tibiae were decalcified in 14% w/v EDTA at 4°C and embedded in paraffin. 5μm slices were stained with hematoxylin and eosin or with tartrate resistant acid phosphatase by the Imaging and Histology Laboratory at the Centre for Bone and Periodontal Research (McGill University, Quebec, Canada). Sections were visualized on an Olympus BX51 light microscope (Olympus America Inc., Center Valley, PA).

### Gene expression analyses

Using femoral bone marrow samples or crushed, whole tibia, total RNA was extracted and genomic DNA was removed using the Direct-Zol™ Miniprep Kit with an in column DNase I treatment (Zymo Research, Irvine, CA). cDNA was prepared from total RNA using the iScript™ Reverse Transcription System (Bio-Rad Laboratories, Inc., Hercules, CA). All qPCR reactions were performed using the PowerUp™ SYBR Green Master Mix (Thermo Fisher). The qPCR reactions were performed using a CFX96 Real-Time System (Bio-Rad) or a QuantStudio3 Real-Time PCR System (Thermo Fisher Scientific, Waltham, MA): hot-start activation at 95°C for 2 min, 40 cycles of denaturation (95°C for 15 sec) and annealing/extension (<60°C for 15 sec and 72°C for 60 sec or ≥60°C for 60 sec). qPCR reactions were run in duplicate. The primer sequences and annealing temperatures are provided in **Table 1**. Relative gene expression was determined using the Pfaffl method to account for differential primer efficiencies [34], using the geometric mean of the Cq values for beta-2-microglobulin (*B2m*) and 18s ribosomal RNA (*Rn18s*) for normalization. The average Cq value from vehicle-treated, sham, low fat mice was used as the reference point. Data are reported as “Relative Expression.”

**Table 1.**
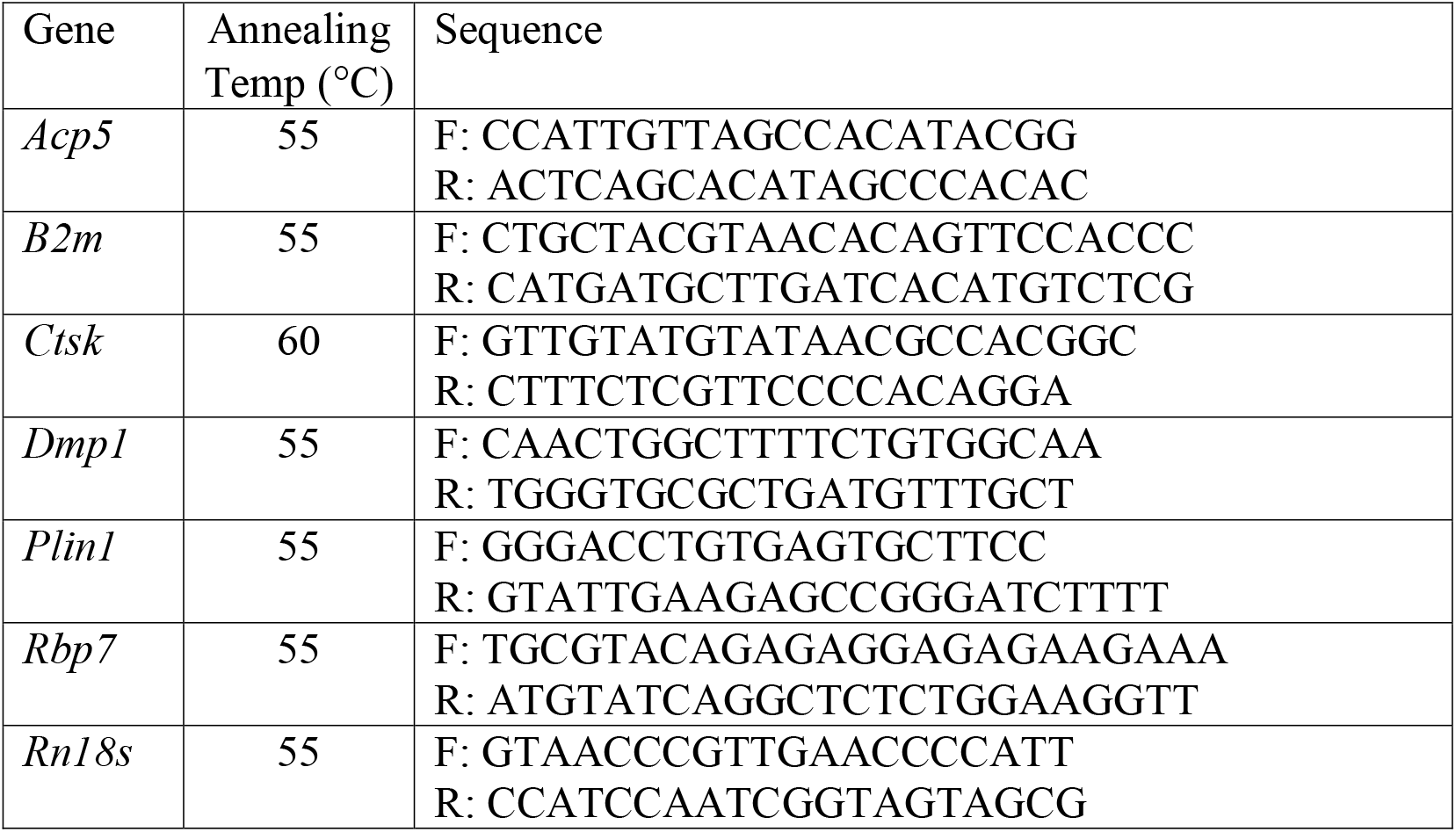
Primers used for qPCR.

### Statistical analyses

Data are presented from individual mice with the mean indicated with a line. A total of 13 outliers were identified as follows: weight gain, liver weight, adipose weight and spleen weight were each averaged, and animals with any value that was 2 SD ± the mean were identified and removed from all analyses. Physiological data from a treatment group are from 6-16 individual mice, depending on the assay. The elemental Sn analyses are from pools of 4 mice each (4 pools per group). Statistical analyses were performed with Prism 8 (GraphPad Software Inc., La Jolla, CA). One-way or two-way ANOVAs with the Dunnett’s or Sidak’s post hoc test were used to determine statistical significance. All analyses were performed using α = 0.05.

## Results

### Metabolic Characteristics

TBT is a known metabolic disruptor [35], therefore we examined several aspects of metabolic homeostasis. Skeletally mature, female mice (12 weeks of age [36]) that had undergone sham surgery or were ovariectomized at 10 weeks of age were treated 3 times/week for 14 weeks with either vehicle (sesame oil) or TBT (1 and 5 mg/kg bw) by oral gavage and fed a low (10%) or high (45%) fat diet. Across all mice, adiposity was highly correlated with the rate of weight gain (Pearson r = 0.9039, p <0.0001, **Figure 1a**). In low fat fed mice, OVX significantly increased adiposity (**Figure 1b**). TBT had no effect on adiposity in intact, low fat fed mice (**Figure 1b**), but TBT suppressed the increase in adiposity in OVX mice (**Figure 1b**). In high fat fed mice, OVX significantly increased adiposity (**Figure 1b**). TBT significantly increased adiposity in intact, high fat fed mice (**Figure 1b**), but TBT modestly, although not significantly, suppressed the increase in adiposity in OVX mice (**Figure 1b**). In mice fed a low-fat diet, only TBT treatment in OVX mice significantly increased lipid accumulation in liver (**Figure 1c**). In mice fed a high fat diet, OVX significantly increased lipid accumulation in liver, while TBT had no significant affect (**Figure 1c**).

**Figure 1.**
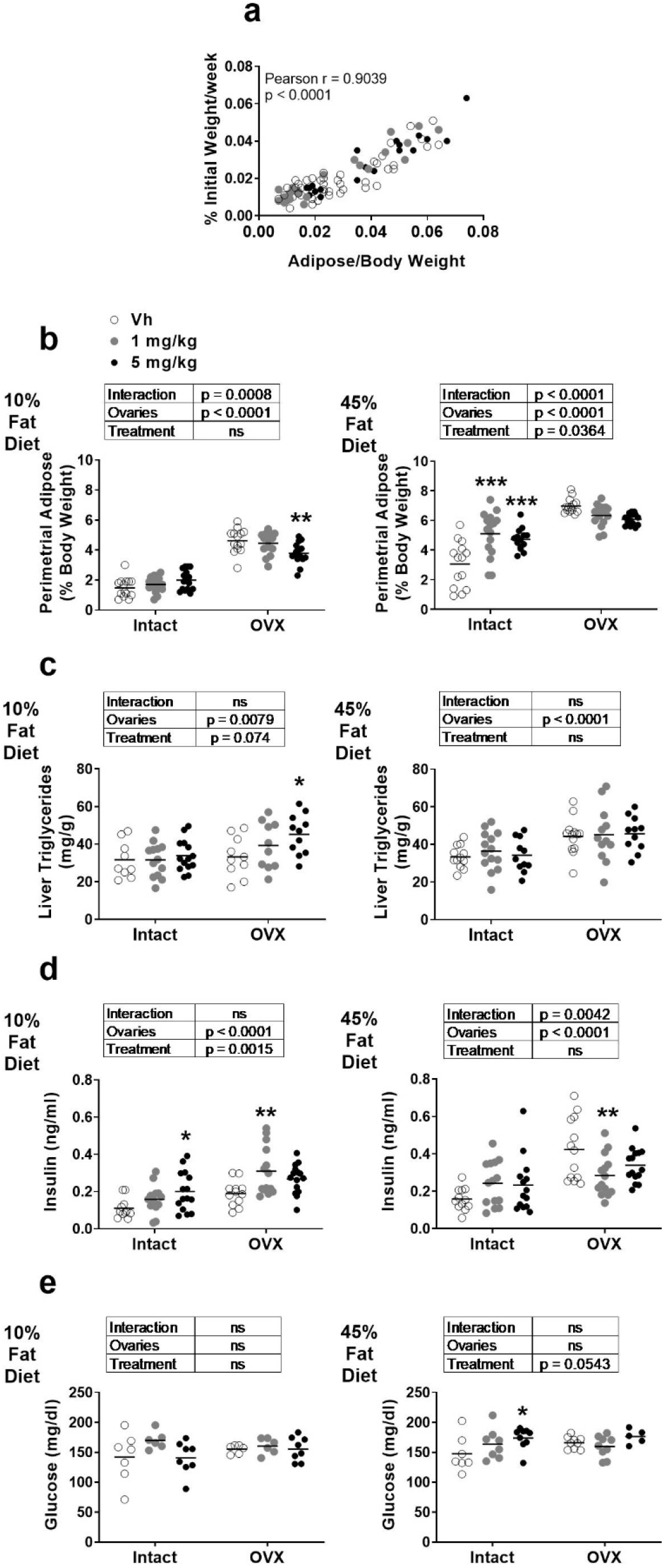
Metabolic parameters in female intact or ovariectomized (OVX) C57Bl/6J mice fed a 10% or 45% fat diet and treated with TBT. Panel **a:** Rate of weight gain versus parametrial adipose weight. Mice across the four cohorts are indicated by shading of the points. Panel **b:** Adipose weight. Panel **c**: Liver triacylglyceride concentration. Panel **d**: Serum insulin. Panel **e:** Serum glucose. Data are presented from individual mice, and the mean is indicated by a line. n=14-16 individual mice. Boxes show results of Two-way ANOVA. *p < 0.05, **p<0.01 versus Vh, Dunnett’s multiple comparison test.

Indicators of glucose homeostasis also were altered. In mice fed a low-fat diet, OVX significantly increased serum insulin (**Figure 1d**). Further, TBT significantly increased circulating insulin in both intact and OVX mice (**Figure 1d**). In mice fed a high-fat diet, OVX significantly increased serum insulin (**Figure 1d**). TBT did not have a significant effect on serum insulin in intact, high fat fed mice, but significantly decreased serum insulin in OVX mice (**Figure 1d**). In low fat fed mice, serum glucose was largely unaffected by OVX or TBT exposure (**Figure 1e**). In high fat fed mice, OVX did not have an effect on circulating glucose (**Figure 1e**). TBT increased serum glucose with a more significant effect in intact than OVX mice (**Figure 1e**).

### Tin (Sn) Concentrations in Bone

In order to determine the amount of TBT reaching the bone, we measured a surrogate of TBT concentration, elemental Sn, in whole humerus. Sn content was increased significantly in bone of TBT treated mice, regardless of diet (**Figure 2**). A significant caveat, however, is that it is unknown what proportion of this is bioavailable TBT. Because the 5 mg/kg dose significantly increased Sn content, we proceeded with analyzing bone structure and gene expression in the groups receiving this dose, in comparison to Vh.

**Figure 2.**
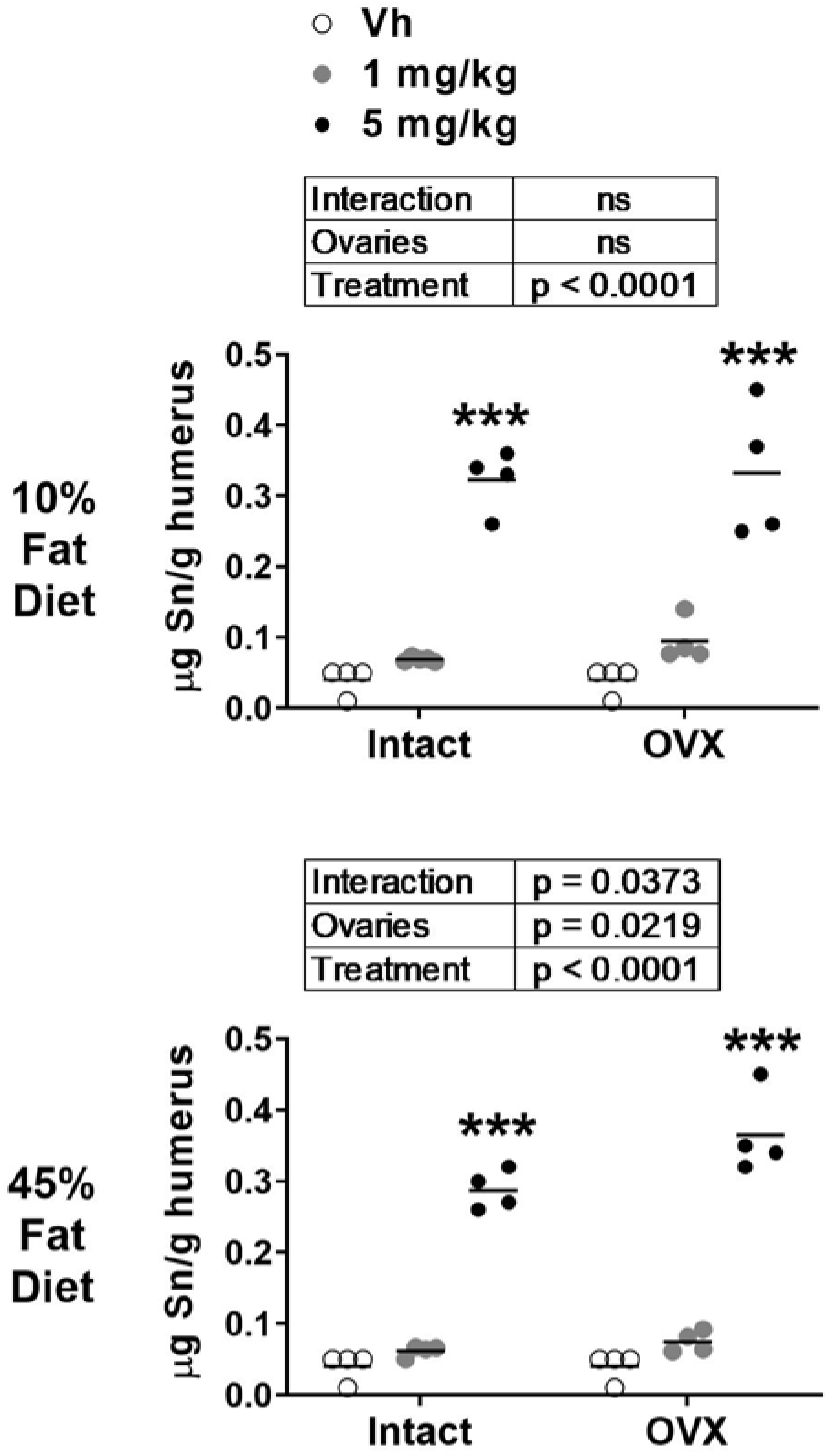
Analysis of elemental tin in bone after TBT treatment. Elemental tin (Sn) content was determined in humeri by ICP-MS. Data are presented from pools of mice (4-5 mice per pool), and the mean is indicated by a line. n=4-5 pool individual mice. Boxes show results of Two-way ANOVA. **p<0.01 vs. Vh, Dunnett’s multiple comparison test.

### Micro-CT Analyses

Micro-CT was used to assess TBT’s impact on the microarchitecture of the femur. The cortical compartment was analyzed at the diaphysis of the femur. In low fat fed, OVX mice, cortical area (Ct.Ar), the ratio of cortical area to total area (Ct.Ar/Tt.Ar), and cortical thickness (Ct.Th) were significantly lower, while marrow area (Ma.Ar) was significantly higher, relative to intact mice. Moment of inertia (MOI) and cortical tissue mineral density were unaffected by OVX (**Figure 3a-b**). TBT treatment in low fat fed mice resulted in significantly lower Tt.Ar and Ma.Ar across intact and OVX mice (**Figure 3a-b**). In high fat fed, OVX mice, Tt.Ar and Ma.Ar were significantly higher, while Ct.Ar and Ct.Ar/Tt.Ar were significantly lower, relative to intact mice (**Figure 3c-d**). Tissue mineral density also was significantly higher in OVX mice fed a high fat diet relative to intact mice (**Figure 3c-d**). TBT treatment in high fat fed mice resulted in a significantly lower Ct.Ar, Ct.Ar/Tt.Ar, Ct.Th and MOI across intact and OVX mice, with TBT inducing a significant decrease in Ct.Ar and Cr.Th in OVX mice relative to Vh-treated mice (**Figure 3c-d**). TBT treatment did not have a significant effect on cortical tissue mineral density in high fat fed, OVX or intact mice (**Figure 3c-d**).

**Figure 3.**
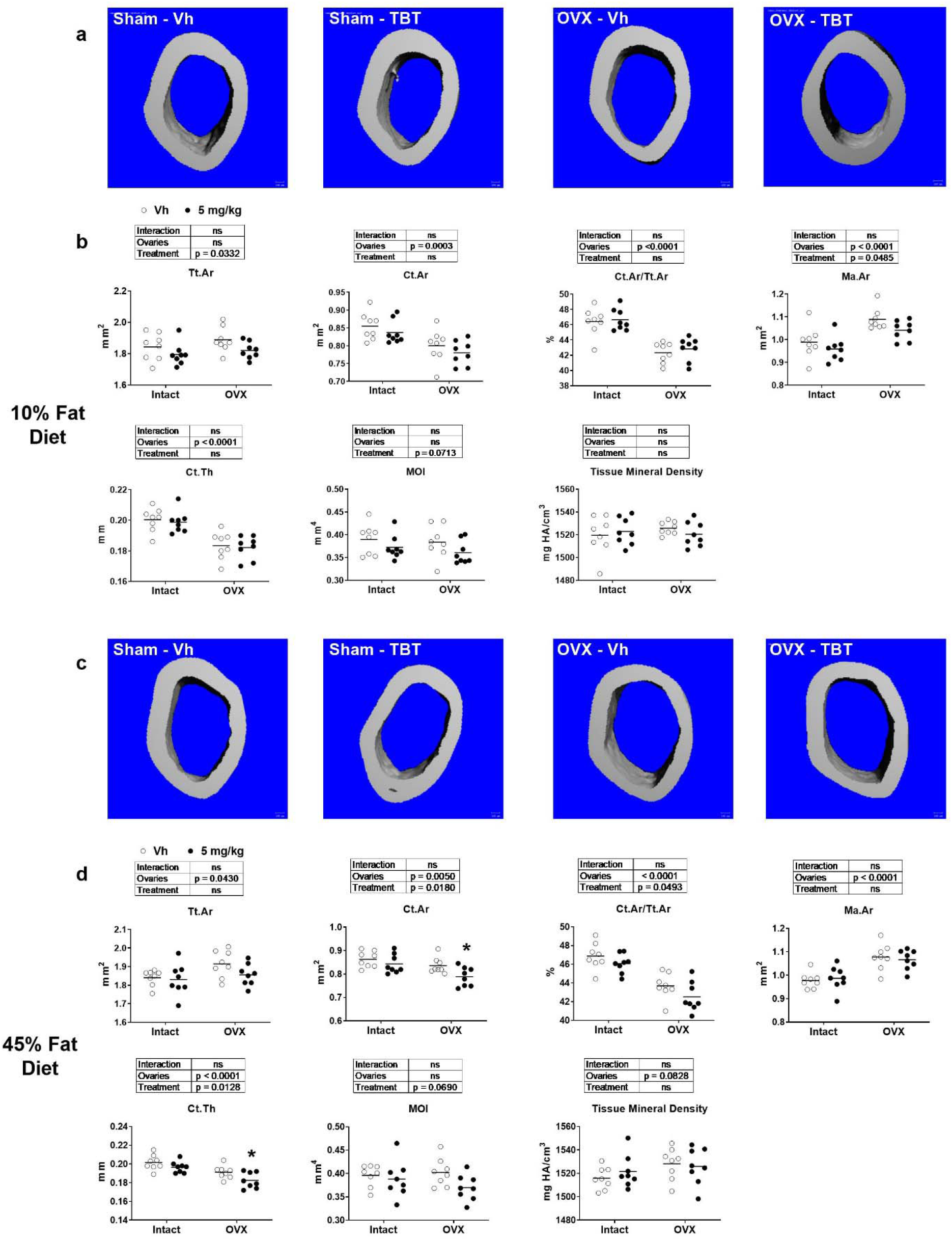
micro-CT analysis of femur cortical parameters. Panel **a**: Representative micro-CT images of mid-diaphysis of low-fat fed mice. Panel **b**: Cortical bone parameters of low-fat fed mice. Panel **c**: Representative micro-CT images of mid-diaphysis of high fat fed mice. Panel **d**: Cortical bone parameters of high fat fed mice. Ct.Th: cortical bone thickness; Ct.Ar: cortical bone area; Ma.Ar: medullary (marrow) area; Tt.Ar: total cross-sectional area; Ct.Ar/Tt.Ar: cortical area fraction; MOI: moment of inertia; mineral density. Data are presented from individual mice, and the mean is indicated by a line. n=6 individual mice. Boxes show results of Two-way ANOVA. *p<0.05, **p<0.01 versus Vh, Sidak’s multiple comparison test.

The trabecular compartment was analyzed at the metaphysis of the femur. In low fat fed, OVX mice, trabecular number (Tb.N) and connective density (Conn.D) were significantly lower and trabecular spacing (Tb.Sp) was significantly higher relative to intact mice (**Figure 4a-b**). The bone volume fraction (BV/TV), trabecular tissue mineral density, structure model index (SMI), and trabecular thickness (Tb.Th) were not different between low fat fed, intact and OVX mice (**Figure 4a-b**). TBT treatment in low fat fed mice resulted in a significantly higher BV/TV, Conn. D and Tb.N and significantly lower Tb.Sp across intact and OVX mice relative to Vh-treated mice, with a significant interaction occurring between ovary status and TBT treatment (**Figure 4a-b**). TBT only significantly reduced trabecular tissue mineral density in intact mice, TBT reduced SMI across intact and OVX mice, and TBT had no effect on Tb.Th in either intact of OVX mice (**Figure 4a-b**). In high fat fed, OVX mice, BV/TV, Conn.D, and Tb.N were significantly lower, and Tb.Sp was significantly higher, relative to intact mice (**Figure 4c-d**). TBT treatment in high fat fed mice resulted in significantly lower SMI and Tb.Sp, as well as significantly higher Conn.D and Tb.N relative to Vh across intact and OVX mice (**Figure 4c-d**). Again, trabecular thickness was not influenced by treatment or ovary status in high fat fed mice.

**Figure 4.**
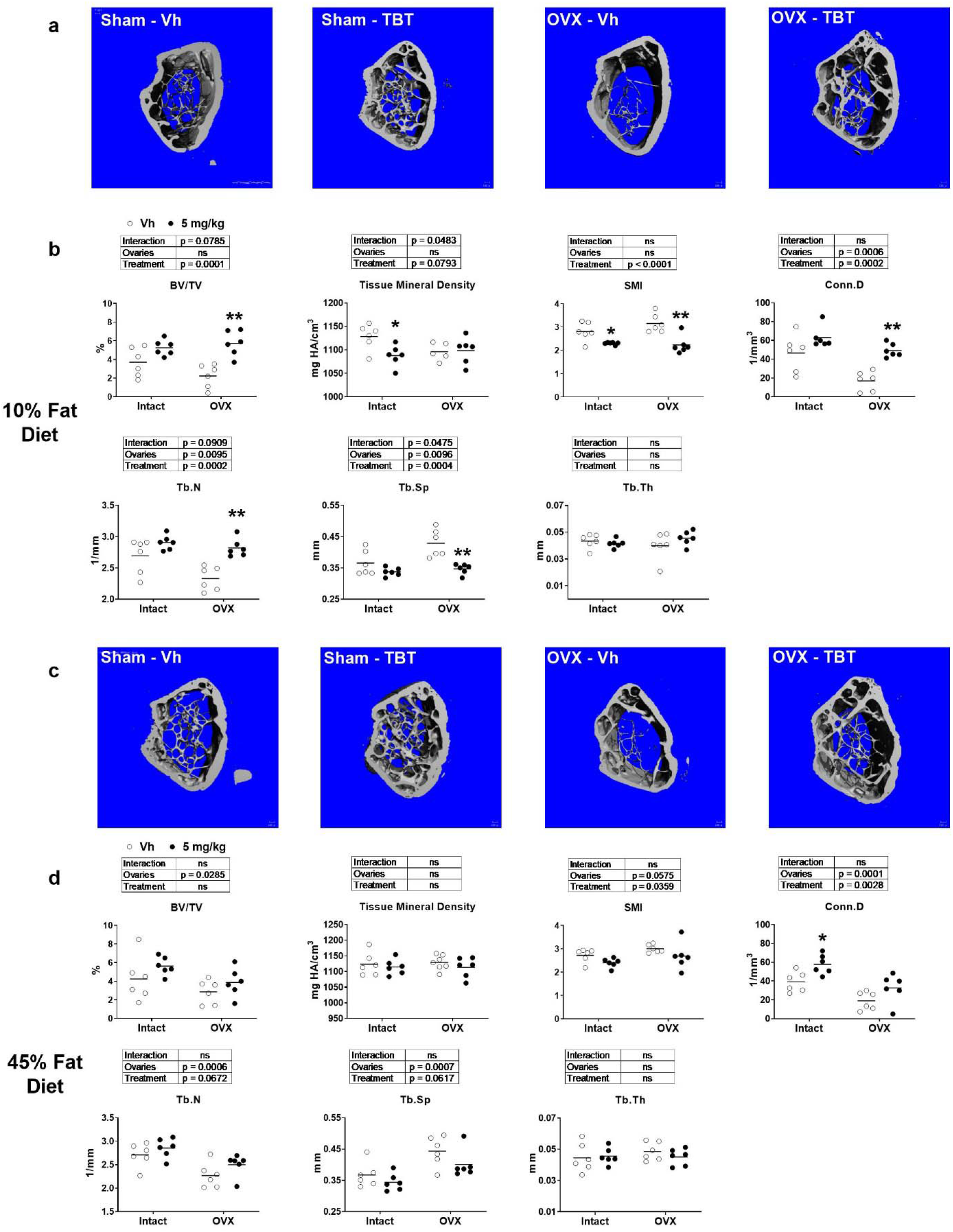
μCT analysis of femur trabecular parameters. Panel **a:** Representative micro-CT images of distal metaphysis of low-fat fed mice Panel **b:** Trabecular bone parameters of low-fat fed mice. Panel **c**: Representative micro-CT images of distal metaphysis of high fat fed mice. Panel **d:** Trabecular bone parameters of high fat fed mice. BV/TV: bone volume fraction; mineral density; SMI: structure model index; Conn.D: connectivity density; Tb.N: trabecular number; Tb.Sp: mean trabecular spacing; Tb.Th: mean trabecular thickness. Data are presented from individual mice, and the mean is indicated by a line. n=6 individual mice. Boxes show results of Two-way ANOVA. *p<0.05, **p<0.01versus Vh, Sidak’s multiple comparison test.

### Histology and Biochemical Analyses

Histology revealed that the most obvious characteristic of bones from TBT-treated, low fat fed mice, both intact and OVX, was the increase in adipocytes in the marrow cavity of the proximal tibia (**Figure 5a**). This was accompanied by a significantly higher expression of the adipocyte-specific gene *Plin1* in whole tibia in intact and OVX mice (**Figure 5b**). The expression of the osteoblast-specific gene *Dmp1* was highly variable and not significantly affected by OVX or TBT in low fat fed mice (**Figure 5b**). The bone formation marker, PINP, was not different in serum of any low-fat fed mice; however, serum osteocalcin was significantly higher in OVX mice (**Figure 5c**). There also was an interaction between ovary status and TBT treatment, with TBT treatment being associated with increasing serum osteocalcin concentrations in intact mice but decreasing serum osteocalcin concentrations in OVX mice (**Figure 5c**). Bone marrow adiposity also was evident in high fat fed intact and OVX mice (**Figure 5d**). TBT treatment significantly increased expression of *Plin1* in whole tibia across intact and OVX mice, and there was no effect on expression of *Dmp1* (**Figure 5e**). Serum PINP and osteocalcin concentrations were decreased most significantly in TBT-treated OVX mice fed a high fat diet (**Figure 5f**).

**Figure 5.**
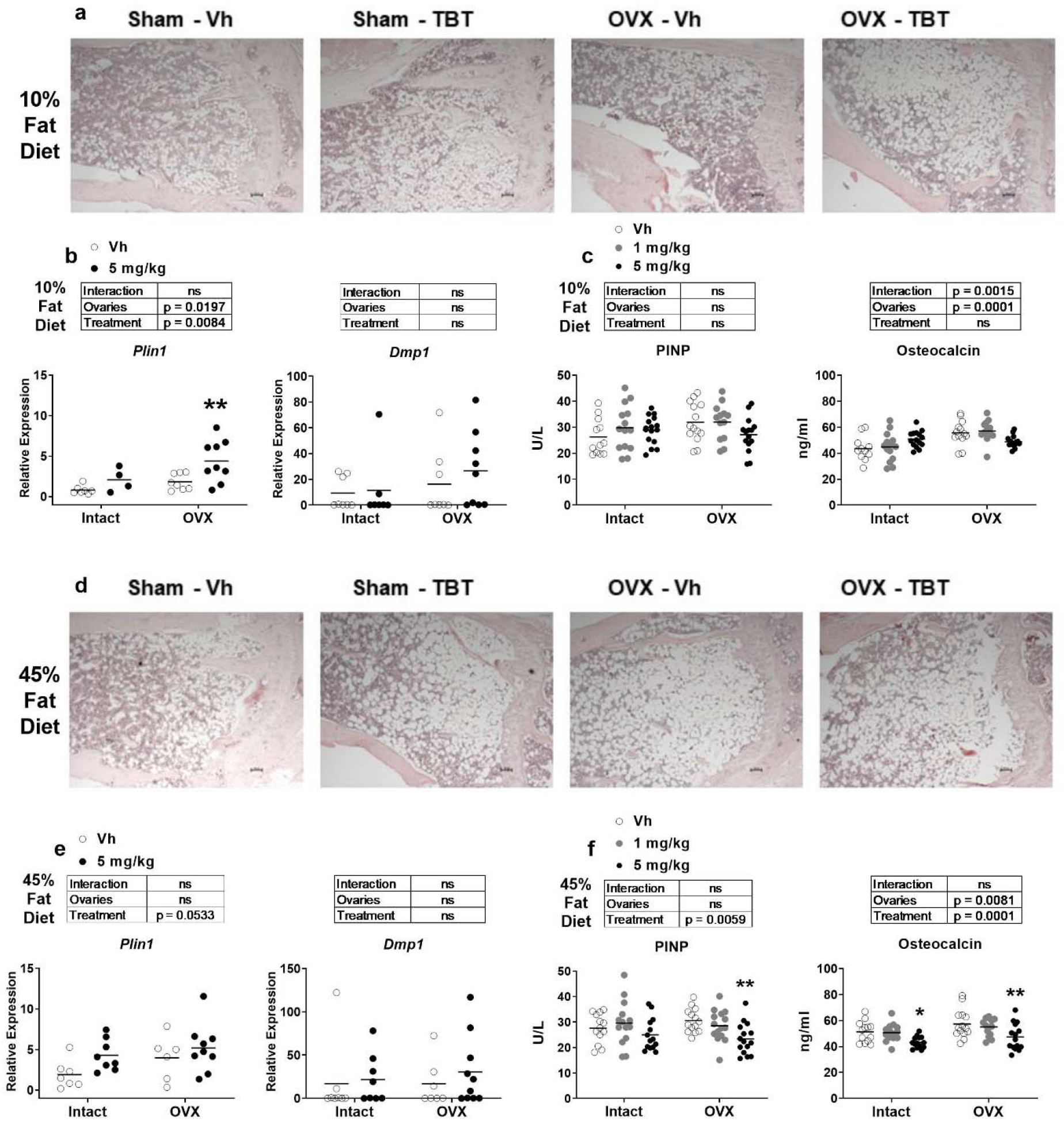
Analyses of bone formation. Panels **a** and **d**: Representative images of 5 μm slices of distal femur (n=4-6) stained with hematoxylin and eosin (4x magnification). Scale bar = 100 μm Panels **b** and **e**: mRNA expression of an adipocyte (*Plin1*) and osteoblast (*Dmp1*) biomarker genes from whole tibia (n=7-9). Panels **c** and **f**: Quantification of serum PINP and OCN (n=11-15). Data are presented from individual mice, and the mean is indicated by a line. Boxes show results of Two-way ANOVA. *p < 0.05, **p<0.01 versus Vh, Sidak’s (c, d) or Dunnett’s (e, f) multiple comparison test.

Osteoclast number and activity also were examined. In low fat fed mice, TRAP staining of the proximal tibia showed little difference in between intact/OVX mice or treatment groups (**Figure 6a**). This was consistent with there being no differences in the expression of *Acp5* or *Ctsk* in bone marrow in either intact or OVX mice (**Figure 6b**). However, serum Trap5b and CTX were significantly lower in TBT-treated, OVX mice (**Figure 6c**). In high fat fed mice, TRAP staining of the proximal tibia also showed little difference in between intact/OVX mice or treatment groups (**Figure 6d**). While TBT had no effect on expression of *Acp5* or *Ctsk* in intact, high fat fed mice, TBT significantly decreased their expression in OVX mice (**Figure 6e**). Serum Trap5b did not differ between intact/OVX mice or treatment groups, but serum CTX was significantly reduced by TBT in high fat fed, OVX mice (**Figure 6f**).

**Figure 6.**
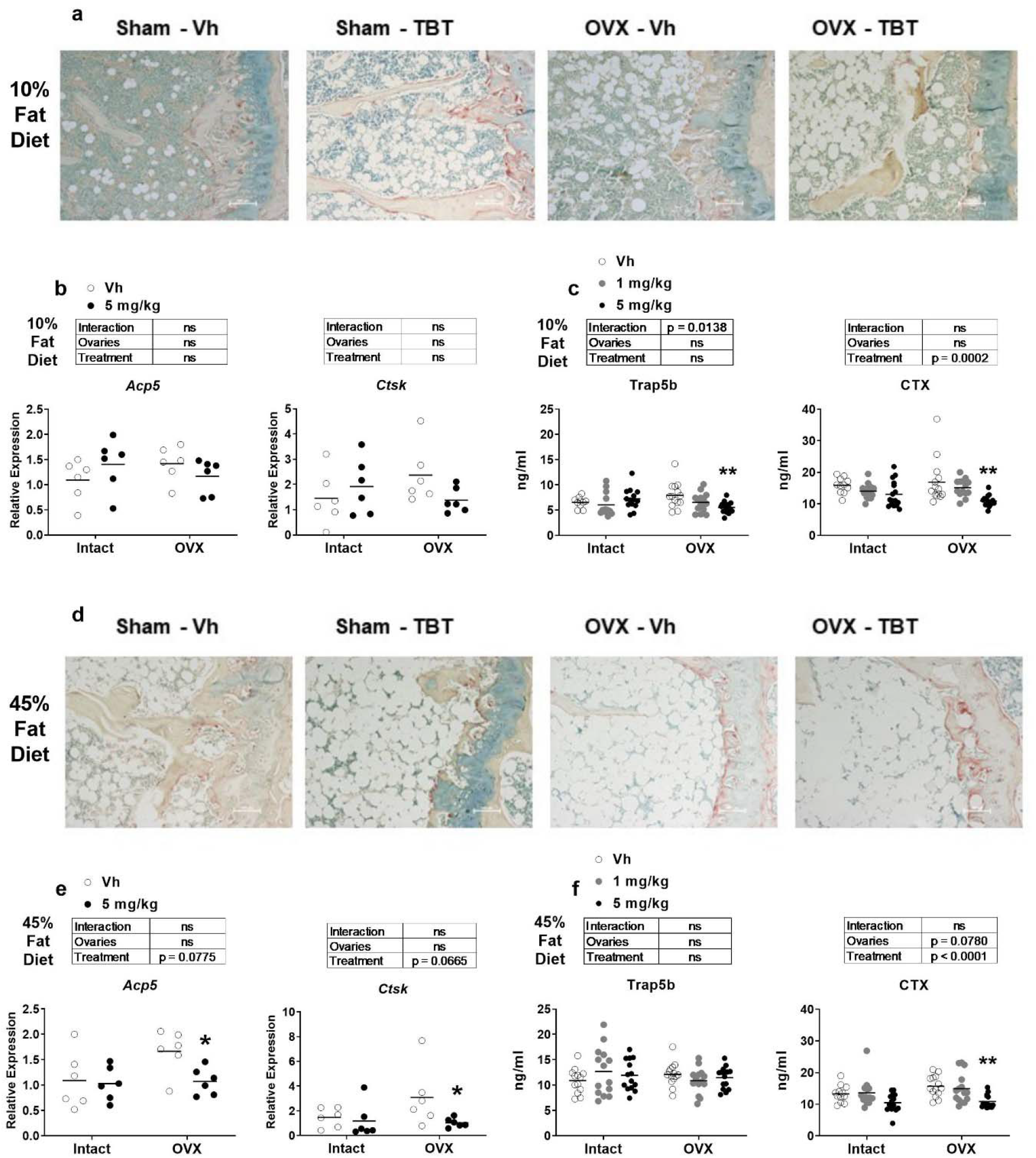
Analyses of bone resorption. Panels **a** and **d**: Representative images of 5 μm slices of distal femur (n=4-6) stained for TRAP (red shading)(10x magnification). Scale bar = 100 μm. Panels **b** and **e**: mRNA expression of osteoclast biomarker genes from femur bone marrow (n=7-9). Panels **c** and **f**: Quantification of serum Trap5b and CTX (n = 11-15). Data are presented from individual mice, and the mean is indicated by a line. Boxes show results of Two-way ANOVA. *p < 0.05, **p<0.01 versus Vh, Sidak’s (c, d) or Dunnett’s (e, f) multiple comparison test.

## Discussion

TBT is a unique nuclear receptor ligand with the ability to directly bind and activate PPARγ and RXR and to indirectly activate LXR [21, 37, 38]. We previously showed that while TBT treatment reduced bone formation in the cortical compartment, consistent with *in vitro* studies of bone forming cells demonstrating that TBT inhibits osteogenesis, it increased trabecular bone likely through modulating osteoclast phenotype [26]. The goal of this study was to investigate the effect of TBT on bone structure in two scenarios associated with bone loss, consumption of a high fat diet and ovariectomy.

Given that Sn concentrations were significantly elevated in bone of TBT-treated mice, it is likely that the bone phenotypes observed are a result of direct action of TBT on bone and bone marrow cells. *In vitro* studies of osteoblasts and osteoclasts in mono-culture have shown that TBT activates PPARγ, RXR and LXR in bone forming cells [26, 37, 39] and RXR and LXR in osteoclast forming cells [26, 38]. TBT skews multipotent stromal cell differentiation away from osteogenesis and towards adipogenesis [26, 37, 39]. In osteoclast-forming cultures, TBT has a minimal effect on the number of osteoclasts that differentiate, but it significantly changes the function of osteoclasts at the transcriptional level decreasing expression of function markers and increasing expression of pro-osteogenic signals and decreasing resorptive activity [26, 29].

TBT had marginal effects on cortical and trabecular bone in intact mice fed a low- or high-fat diet. Across intact and OVX mice there was a reduction in total cortical area and in marrow area in the low-fat fed mice but not the high-fat fed mice. We observed these changes in our previous study and hypothesized these changes resulted from a failure of periosteal bone deposition accompanied by a decrease in osteoclast activity at the endosteal surface [26]. We did not observe TBT-mediated thinning of the cortex at mid-diaphysis or enhancement of trabecular bone volume (increased trabecular number without an increase in average trabecular thickness) in TBT-treated mice. This difference from our previous observations [26] may most likely be because the doses used in the current study were ten- and two-fold lower than in the previous study. Consistent with a lower dose are the observations that TBT treatment had minimal effects on expression of osteoblast or osteoclast biomarker genes and serum makers of bone formation or resorption.

OVX resulted in significant effects on cortical and trabecular bone including reduced bone area, bone area:total area, and cortical thickness, as well as an increase in marrow area, regardless of diet. These observations are largely consistent with those previously reported by Bouxsein et al [40]. The differences (e.g., lack of reduction of total area and significant increase in marrow area) may likely be explained by the fact that in the study by Bouxsein et. al the mice were examined 4 weeks after surgery, while in this study mice were examined 14 weeks after surgery. OVX decreased trabecular number and connective density, while increasing trabecular spacing, regardless of diet. Additionally, OVX only decreased trabecular bone volume fraction in mice fed a high fat diet. Again, these observations are largely consistent with those previously reported [40]; the difference (e.g., significant increase in trabecular spacing and significant decrease in bone volume fraction only in high fat diet mice) may be attributed to differences in anatomical site. The study by Bouxsein et al examined trabecular bone in the proximal tibia, while in this study trabecular bone was examined in the distal femur.

TBT had a significant effect on cortical and trabecular bone in OVX mice. In high-fat fed OVX mice, TBT modestly exacerbated OVX-mediated reductions in cortical thickness, bone area and total area. In contrast, TBT significantly (i.e., there was a significant interaction) mitigated OVX-mediated reductions in trabecular bone volume fraction, trabecular number and trabecular spacing, along with a modest mitigation of OVX-mediated reduction in connective density. These results and the observation that TBT induced an increase in whole body bone mineral density [27] provide independent corroboration of our previous observation that TBT treatment can increase trabecular bone.

TBT did not significantly protect mice from OVX-induced reductions in trabecular bone in high fat fed mice, however. The loss of protective affect was accompanied by significant decreases in serum bone formation markers, which appear to have outweighed the TBT-induced reduction in osteoclast activity, indicated by the decreased expression of *Ctsk* in bone marrow and reduced serum CTX. The change in the balance of TBT’s effects on osteoblasts and osteoclasts may result from a change in expression of PPARγ, the expression of which is increased by a high fat diet [15, 16]. This is supported by the fact that adiposity was significantly increased in the bone marrow of OVX mice by TBT, along with the expression of the adipocyte-specific, PPARγ target gene *Plin1*.

TBT is a toxic agent and is well known as an immunotoxicant [41]. In this study, the 5 mg/kg dose of TBT reduced absolute spleen weight in all intact mice fed a high fat diet (not shown). Additionally, TBT is a metabolic disruptor [35], which has been shown to increase adiposity and disrupt glucose homeostasis [42]. We observed that TBT significantly increased weight gain, which was highly correlated with an increase in adiposity. Furthermore, TBT increased fasting insulin in low fat fed, intact mice, with a similar trend in high fat fed, intact mice. TBT also increased fasting glucose in high fat fed, intact mice. In contrast, TBT mitigated OVX-induced weight gain and adiposity and reduced serum insulin levels. Classically, TBT is thought of as an aromatase inhibitor [35]; however, evidence also suggests that TBT can enhance estradiol synthesis [43, 44]. A potential mechanism by which TBT could mitigate the effects of OVX could be through stimulating estrogen production by extra-gonadal tissues [45].

The effects of TBT on bone make it clear that its complex activation of PPARγ, RXR and LXR make it unsuitable as a model for therapeutics. However, the results of this study do support the potential for targeting LXR and/or RXR in interventions to combat low bone density, given their roles in moderating osteoclast differentiation [30, 31]. The results also show that diet is likely to significantly confound the efficacy of intervention designed to slow osteoporosis. Future studies should be conducted to tailor the effect of LXR/RXR ligands to selectively activate pathways in bone cells.

## Acknowledgments

This work was supported by the National Institute of Environmental Health Sciences grant R21 ES021136. The authors thank Ms. Cassie Huang for her excellent technical assistance.

## References

1. Rosen CJ, Bouxsein ML (2006) Mechanisms of disease: is osteoporosis the obesity of bone? Nat Clin Pr Rheumatol 2:35–43

2. Moerman EJ, Teng K, Lipschitz DA, Lecka-Czernik B (2004) Aging activates adipogenic and suppresses osteogenic programs in mesenchymal marrow stroma/stem cells: the role of PPAR-gamma2 transcription factor and TGF-beta/BMP signaling pathways. Aging Cell 3:379–389

3. Akune T, Ohba S, Kamekura S, et al (2004) PPARgamma insufficiency enhances osteogenesis through osteoblast formation from bone marrow progenitors. J Clin Invest 113:846–855

4. Lecka-Czernik B, Gubrij I, Moerman EJ, et al (1999) Inhibition of Osf2/Cbfa1 expression and terminal osteoblast differentiation by PPARgamma2. J Cell Biochem 74:357–371

5. Baroi S, Czernik PJ, Chougule A, et al (2021) PPARG in osteocytes controls sclerostin expression, bone mass, marrow adiposity and mediates TZD-induced bone loss. Bone 115913. https://doi.org/10.1016/j.bone.2021.115913

6. Wan Y, Chong LW, Evans RM (2007) PPAR-gamma regulates osteoclastogenesis in mice. Nat Med 13:1496–1503

7. Wei W, Wang X, Yang M, et al (2010) PGC1beta mediates PPARgamma activation of osteoclastogenesis and rosiglitazone-induced bone loss. Cell Metab 11:503–516

8. Billington EO, Grey A, Bolland MJ (2015) The effect of thiazolidinediones on bone mineral density and bone turnover: systematic review and meta-analysis. Diabetologia 58:2238–2246. https://doi.org/10.1007/s00125-015-3660-2

9. Bilik D, McEwen LN, Brown MB, et al (2010) Thiazolidinediones and fractures: evidence from translating research into action for diabetes. J Clin Endocrinol Metab 95:4560–4565

10. Dong X-L, Li C-M, Cao S-S, et al (2016) A High-Saturated-Fat, High-Sucrose Diet Aggravates Bone Loss in Ovariectomized Female Rats. J Nutr 146:1172–1179. https://doi.org/10.3945/jn.115.225474

11. Wronski TJ, Cintron M, Doherty AL, Dann LM (1988) Estrogen treatment prevents osteopenia and depresses bone turnover in ovariectomized rats. Endocrinology 123:681–686

12. Melville KM, Kelly NH, Khan SA, et al (2014) Female mice lacking estrogen receptor-alpha in osteoblasts have compromised bone mass and strength. J Bone Min Res 29:370–379

13. Sottile V, Seuwen K, Kneissel M (2004) Enhanced marrow adipogenesis and bone resorption in estrogen-deprived rats treated with the PPARgamma agonist BRL49653 (rosiglitazone). Calcif Tissue Int 75:329–337

14. Cho SW, Yang JY, Her SJ, et al (2011) Osteoblast-targeted overexpression of PPARgamma inhibited bone mass gain in male mice and accelerated ovariectomy-induced bone loss in female mice. J Bone Min Res

15. Cao JJ, Gregoire BR, Gao H (2009) High-fat diet decreases cancellous bone mass but has no effect on cortical bone mass in the tibia in mice. Bone 44:1097–1104

16. Chen JR, Lazarenko OP, Wu X, et al (2010) Obesity reduces bone density associated with activation of PPARgamma and suppression of Wnt/beta-catenin in rapidly growing male rats. PLoS One 5:e13704

17. Sherk VD, Heveran CM, Foright RM, et al (2021) Sex differences in the effect of diet, obesity, and exercise on bone quality and fracture toughness. Bone 145:115840. https://doi.org/10.1016/j.bone.2021.115840

18. Halade G V, Rahman MM, Williams PJ, Fernandes G (2010) High fat diet-induced animal model of age-associated obesity and osteoporosis. J Nutr Biochem 21:1162–1169

19. Patsch JM, Kiefer FW, Varga P, et al (2011) Increased bone resorption and impaired bone microarchitecture in short-term and extended high-fat diet-induced obesity. Metabolism 60:243–249

20. Kanayama T, Kobayashi N, Mamiya S, et al (2005) Organotin compounds promote adipocyte differentiation as agonists of the peroxisome proliferator-activated receptor gamma/retinoid X receptor pathway. Mol Pharmacol 67:766–774

21. le Maire A, Grimaldi M, Roecklin D, et al (2009) Activation of RXR-PPAR heterodimers by organotin environmental endocrine disruptors. EMBO Rep 10:367–373

22. Baker AH, Wu TH, Bolt AM, et al (2017) Tributyltin alters the bone marrow microenvironment and suppresses B cell development. Toxicol Sci 158:. https://doi.org/10.1093/toxsci/kfx067

23. Watt J, Schlezinger JJ (2015) Structurally-diverse, PPARγ-activating environmental toxicants induce adipogenesis and suppress osteogenesis in bone marrow mesenchymal stromal cells. Toxicology 331:. https://doi.org/10.1016/j.tox.2015.03.006

24. Koskela A, Viluksela M, Keinänen M, et al (2012) Synergistic effects of tributyltin and 2,3,7,8-tetrachlorodibenzo-p-dioxin on differentiating osteoblasts and osteoclasts. Toxicol Appl Pharmacol 263:210–217. https://doi.org/10.1016/j.taap.2012.06.011

25. Tsukamoto Y, Ishihara Y, Miyagawa-Tomita S, Hagiwara H (2004) Inhibition of ossification in vivo and differentiation of osteoblasts in vitro by tributyltin. Biochem Pharmacol 68:739–746

26. Watt J, Baker AH, Meeks B, et al (2018) Tributyltin induces distinct effects on cortical and trabecular bone in female C57Bl/6J mice. J Cell Physiol 233:. https://doi.org/10.1002/jcp.26495

27. Resgala LCR, Santana HS, Portela BSM, et al (2019) Effects of Tributyltin (TBT) on Rat Bone and Mineral Metabolism. Cell Physiol Biochem Int J Exp Cell Physiol Biochem Pharmacol 52:1166–1177. https://doi.org/10.33594/000000079

28. Yao W, Wei X, Guo H, et al (2020) Tributyltin reduces bone mineral density by reprograming bone marrow mesenchymal stem cells in rat. Environ Toxicol Pharmacol 73:103271. https://doi.org/10.1016/j.etap.2019.103271

29. Yonezawa T, Hasegawa S, Ahn JY, et al (2007) Tributyltin and triphenyltin inhibit osteoclast differentiation through a retinoic acid receptor-dependent signaling pathway. Biochem Biophys Res Commun 355:10–15. https://doi.org/10.1016/j.bbrc.2006.12.237

30. Goel D, Vohora D (2021) Liver X receptors and skeleton: Current state-of-knowledge. Bone 144:115807. https://doi.org/10.1016/j.bone.2020.115807

31. Menéndez-Gutiérrez MP, Ricote M (2017) The multi-faceted role of retinoid X receptor in bone remodeling. Cell Mol Life Sci 74:2135–2149. https://doi.org/10.1007/s00018-017-2458-4

32. Norris AW, Chen L, Fisher SJ, et al (2003) Muscle-specific PPARgamma-deficient mice develop increased adiposity and insulin resistance but respond to thiazolidinediones. J Clin Invest 112:608–618. https://doi.org/10.1172/JCI17305

33. Bouxsein ML, Boyd SK, Christiansen BA, et al (2010) Guidelines for assessment of bone microstructure in rodents using micro-computed tomography. J Bone Min Res 25:1468–1486

34. Pfaffl MW (2001) A new mathematical model for relative quantification in real-time RT-PCR. Nucleic Acids Res 29:e45

35. Grun F, Watanabe H, Zamanian Z, et al (2006) Endocrine-disrupting organotin compounds are potent inducers of adipogenesis in vertebrates. Mol Endocrinol 20:2141–2155

36. Beamer WG, Donahue LR, Rosen CJ, Baylink DJ (1996) Genetic variability in adult bone density among inbred strains of mice. Bone 18:397–403

37. Baker AH, Watt J, Huang CK, et al (2015) Tributyltin Engages Multiple Nuclear Receptor Pathways and Suppresses Osteogenesis in Bone Marrow Multipotent Stromal Cells. Chem Res Toxicol 28:. https://doi.org/10.1021/tx500433r

38. Cui H, Okuhira K, Ohoka N, et al (2011) Tributyltin chloride induces ABCA1 expression and apolipoprotein A-I-mediated cellular cholesterol efflux by activating LXRα/RXR. Biochem Pharmacol 81:819–824. https://doi.org/10.1016/j.bcp.2010.12.023

39. Kirchner S, Kieu T, Chow C, et al (2010) Prenatal exposure to the environmental obesogen tributyltin predisposes multipotent stem cells to become adipocytes. Mol Endocrinol 24:526–539. https://doi.org/10.1210/me.2009-0261

40. Bouxsein ML, Myers KS, Shultz KL, et al (2005) Ovariectomy-induced bone loss varies among inbred strains of mice. J Bone Min Res 20:1085–1092

41. Baker AH, Wu TH, Bolt AM, et al (2017) From the Cover: Tributyltin Alters the Bone Marrow Microenvironment and Suppresses B Cell Development. Toxicol Sci 158:63–75. https://doi.org/10.1093/toxsci/kfx067

42. Zhan J, Ma X, Liu D, et al (2020) Gut microbiome alterations induced by tributyltin exposure are associated with increased body weight, impaired glucose and insulin homeostasis and endocrine disruption in mice. Environ Pollut 266:115276. https://doi.org/10.1016/j.envpol.2020.115276

43. Nakanishi T, Hiromori Y, Yokoyama H, et al (2006) Organotin compounds enhance 17beta-hydroxysteroid dehydrogenase type I activity in human choriocarcinoma JAr cells: potential promotion of 17beta-estradiol biosynthesis in human placenta. Biochem Pharmacol 71:1349–1357. https://doi.org/10.1016/j.bcp.2006.01.014

44. Munetsuna E, Hattori M, Yamazaki T (2014) Stimulation of estradiol biosynthesis by tributyltin in rat hippocampal slices. Endocr Res 39:168–172. https://doi.org/10.3109/07435800.2013.875563

45. Barakat R, Oakley O, Kim H, et al (2016) Extra-gonadal sites of estrogen biosynthesis and function. BMB Rep 49:488–496. https://doi.org/10.5483/bmbrep.2016.49.9.141

